# Intensity-based axial localization approaches for multifocal plane microscopy

**DOI:** 10.1101/093013

**Authors:** Ramraj Velmurugan, Jerry Chao, Sripad Ram, E. Sally Ward, Raimund J. Ober

**Affiliations:** Department of Molecular and Cellular Medicine, Texas A&M University Health Science Center, College Station, TX 77843, USA; Department of Microbial Pathogenesis and Immunology, Texas A&M University Health Science Center, Bryan, TX 77807, USA; Biomedical Engineering Graduate Program, University of Texas Southwestern Medical Center, Dallas, TX 75390, USA; Department of Biomedical Engineering, Texas A&M University, College Station, TX 77843, USA; Current address: Pfizer, Inc., San Diego, CA, USA

**Keywords:** (180.0180) Microscopy, (100.0100) Image processing, (100.6640) Superresolution, (100.6890) Threedimensional image processing, (100.2550) Focal-plane-array image processors, (180.2520) Fluorescence microscopy

## Abstract

Multifocal plane microscopy (MUM) can be used to visualize biological samples in three dimensions over large axial depths and provides for the high axial localization accuracy that is needed in applications such as the three-dimensional tracking of single particles and superresolution microscopy. This report analyzes the performance of intensity-based axial localization approaches as applied to MUM data using Fisher information calculations. In addition, a new non-parametric intensity-based axial location estimation method, Multi-Intensity Lookup Algorithm (MILA), is introduced that, unlike typical intensity-based methods that make use of a single intensity value per data image, utilizes multiple intensity values per data image in determining the axial location of a point source. MILA is shown to be robust against potential bias induced by differences in the sub-pixel location of the imaged point source. The method's effectiveness on experimental data is also evaluated.

## 1. Introduction

The ability to accurately estimate the three-dimensional (3D) location of a point source can now be achieved through several techniques such as MUltifocal plane Microscopy (MUM) [1–3], the induction of astigmatism in the Point-Spread Function (PSF) [4], and the engineering of the PSF to encode axial information [5, 6]. While all these methods achieve comparable levels of axial localization accuracy, the range of axial locations imaged by a MUM configuration can be increased just by adding more cameras to the setup. While other 3D estimation methods can typically cover axial depths of up to 3 *μ*m [6], MUM configurations with four focal planes can, for example, be used to image through axial depths greater than 8 *μ*m, an imaging window that can cover the full thickness of a typical mammalian cell [7]. To estimate the axial location of a point source using MUM, images of the point source from different detectors, each capturing a different focal plane within the sample, can be fitted to a model that recapitulates the PSF of the microscope configuration. Such a parametric fitting algorithm, named the MUM Localization Algorithm (MUMLA), was first described for the case of MUM with two focal planes [2,8], but has also been extended to MUM with more focal planes [7].

We have also previously developed a Practical Localization Accuracy Measure (PLAM) [9], a quantity based on Fisher information that gives the best accuracy, in terms of the standard deviation, with which a positional coordinate of an object can be estimated given the exact properties of the optical configuration. MUMLA has previously been shown to provide estimates whose standard deviation approaches the PLAM, and hence would be the method of choice if estimates of the highest accuracy are desired. However, a non-parametric algorithm that can rapidly provide an estimate of the axial location of a point source can complement MUMLA. A primary reason is that the fitting of a PSF model to MUM data requires complex PSF fitting algorithms, computational power to perform the fitting, and an appropriate choice of the PSF model. The ability to quickly calculate an estimate of the axial location of a point source can hence allow faster initial analyses of experimental data. Should more accurate results be required, non-parametric estimates can also serve as initial conditions for more rigorous fitting algorithms like MUMLA.

Watanabe and colleagues [10] have previously published such a non-parametric method based on intensity ratios (the ratiometric method) that can be used to quickly calculate the axial location of a point source using data from a two-plane MUM setup. In this method, the peak intensities of a point source in the images from the two focal planes are used to calculate a ratio as described before [1]. Using a pre-determined lookup-table of such ratios for known axial positions of a calibration point source, the axial location of the point source can be estimated based on its calculated ratio.

A similar method that utilizes a sharpness value has been characterized by Greenaway and colleagues [11], where data from MUM configurations with more than two focal planes can also be utilized. These and earlier methods [1] estimate the axial location of a point source using a single intensity metric calculated per focal-plane image of the point source. This motivates the question of how accurately the axial location of a point source can be estimated from such single-intensity calculations, and whether the accuracy of these methods can be improved by modifying these calculations.

We analyze this question by investigating the Fisher information and the associated PLAM that can potentially be attained by using intensity values from different regions of the detectors. These analyses show that at least in some cases, using a single integrated intensity value per detector can provide an axial localization accuracy that is comparable to the accuracy obtained by applying MUMLA on the actual pixelated image as is. However, the PLAM calculations also show that such single-intensity value methods cannot provide consistently good accuracy over large axial ranges.

To address this important drawback, we propose a new axial location estimation approach, Multi-Intensity Lookup Algorithm (MILA), that is based on utilizing intensity values from several different parts of the point source's image. We show that this approach significantly improves on the associated PLAM over the use of a single intensity per detector. We establish that MILA provides good axial localization accuracy over a much wider axial range compared to single-intensity localization methods. We also characterize the bias encountered in non-parametric axial estimation methods due to sub-pixel location differences between point sources and evaluate the ability of MILA to avoid this bias. We evaluate the performance of MILA using both simulated and experimental data. We believe that the versatility and simplicity of our new method will prove useful in the analysis of MUM data.

## 2. Methods

### 2.1. Data simulation

For simulations, images of a point source were modeled as arising from a series of pixelated detectors collecting photons from the point source through a MUM setup. In most cases the Region Of Interest (ROI) was assumed to be a sub-region in the detector with the point source situated such that its location in the image is within the center-pixel of the sub-region. Unless specified otherwise, the point source location is also centered within this pixel. The number of photons detected in each pixel of an ROI is then simulated using the Born and Wolf model for a 3D PSF [12]. A Gaussian random variable with a mean of 80 electrons and a standard deviation of 6 electrons was added to simulate the offset and read-out noise associated with real detectors. A background noise component of 70 photons/pixel/second was also added to all simulations. Simulation of images for a given detector, which captures a distinct focal plane, was carried out with the axial location parameter of the point source offset by the distance of the detector's focal plane from the design location. The photon-detection rate was scaled according to the number of detectors (e.g., if we simulate data as arising from a four-plane MUM setup, a point source from which 8000 photons/s are detected would be simulated such that 2000 photons/s are detected by each of the four cameras). All data were simulated with a wavelength of 655 nm and an exposure time setting of 1 second.

For each dataset, the accuracy benchmark given by the standard PLAM for the z position of the point source was calculated using the FandPLimitTool [2,8,9]. The standard PLAM assumes an estimator that uses each image as is, meaning that as many intensity values as there are pixels in the ROI are used in the estimation. It is the PLAM that has been observed to be approached by the standard deviation of the parametric fitting algorithm MUMLA. In contrast, a Summed-Pixel-Based PLAM (SPB-PLAM) for each dataset was calculated by assuming an estimator that considers a combination of subsets of the ROI, or "sub-ROIs", each represented by a single intensity value obtained as the sum of the intensities of its component pixels. An SPB-PLAM is therefore applicable to intensity-based estimation algorithms such as the sharpness method and MILA. All simulations, PLAM calculations and estimations were performed using code written in MATLAB (MathWorks, Natick, MA), utilizing the EstimationTool [13] and the FandPLimitTool [9,14].

### 2.2. Generation of calibration images from fluorescent beads

To test MILA on experimental data, 100-nm Tetraspeck fluorescent beads (Invitrogen, Carlsbad, CA) were imaged using a four-plane MUM setup. The beads were deposited on MatTek dishes (MatTek, Ashland, MA) pretreated with poly-L-Lysine (Sigma Aldrich) by adding a solution containing the beads for 10 minutes and then washing with water. The dish was then filled with water and imaged using a Zeiss 63× 1.4 NA Plan-Apochromat objective mounted on a piezo nanopositioner (Physik Instrumente, Auburn, MA). Excitation light from a 150-mW, 635-nm solid state laser (OptoEngine, Midvale, UT) was reflected to the objective and the emission light filtered using a quad-band dichroic/emission filter combination (Di01-R405/488/543/635 and FF01-446/515/588/700-25, Semrock, Rochester, NY). The configuration was housed on an Axio Observer.A1 microscope body (Carl Zeiss MicroImaging, Germany). The four-plane MUM configuration was assembled as described previously [2] using two Zeiss dual camera adapters attached on the output ports of another dual camera adapter that is attached to the output port of the microscope body. Four Andor iXon electron-multiplying charge-coupled device (EMCCD) cameras (three iXon DV887 and one iXon DU897, Andor Technology, South Windsor, CT) were attached to the four output ports. The cameras were used to image different focal planes by modifying the length of the spacer between the output port and each camera. Beamsplitters (50/50; Chroma Technology, Bellows Falls, VT) were used to split the incoming emission light to all four cameras. Data was acquired using the conventional read-out mode in all four EMCCD cameras. The lasers, shutters and cameras were synchronously controlled via TTL pulses with custom software written using LabWindows CVI (National Instruments, Austin, TX).

A field of view containing a sparse distribution of beads was selected and z-stack images of this field of view were acquired using the piezo nanopositioner. The images from the different cameras were registered as follows: the lateral coordinates of three or more beads were estimated from the in-focus images of the beads from all the four detectors. The coordinates from the first detector and each of the other three detectors were used to generate affine transformation matrices. The transformation matrices were then used to register the images from the three other detectors to the coordinates of the pixels in the first detector.

To generate the lookup table for MILA, ROIs of selected beads were extracted from each image as small sub-regions such that the center of the bead coincided with the center pixel of the extracted ROI. Such extracted ROIs from all four images per z-slice acquisition of each bead were processed for axial location estimation using MILA.

### 2.3. Identification of the pixel containing the point source location in the image

For the MILA approach, we identify the pixel where the point source is located and calculate the sum intensities of different sub-ROIs around this pixel. In order to efficiently estimate the axial location using this method, the identification of this pixel needs to be performed accurately. This was achieved by fitting a 2D Gaussian profile to the image of the point source. In MUM data where the image of the point source is identifiable in multiple detectors, the image from the detector where the point source was closest to being in focus (which was determined by choosing the image where the most number of photons were detected within the ROI) was used for this estimation.

### 2.4. Estimation of the axial location by MILA

MILA is implemented as follows: The pixel that contains the point source of interest is identified using the images from the in-focus detector in a MUM configuration. Pixels corresponding to concentric square-shaped regions around this center-pixel are then taken from each of these images (Fig. 2A), and their intensities are summed after background subtraction. Hence, if we choose to extract summed intensities from 3 concentric regions in images from a two-plane MUM configuration, we will have 6 intensity values. These intensity values are then normalized by dividing with the highest intensity value among them. Typical normalized intensities from such simulated data are shown as a function of axial position in Fig. 2B. A lookup table of such normalized intensity-values is calculated from images of several point sources at a series of known axial positions. Normalized intensities are then similarly calculated for an unknown point source and compared to the lookup table using a least squares method to estimate the axial location of the point source (a formal description of the method is provided in Appendix 1). For comparison with MILA, the ratiometric method was implemented as described previously [10].

## 3. Results

### 3.1. Comparison of axial localization accuracy between single-intensity methods and MUMLA

Both the ratiometric [10] and the sharpness [11] methods estimate the axial location of the point source by utilizing just a single intensity value from the image of the point source in each detector. In order to understand whether a single integrated intensity value per detector can represent meaningful axial location information, we calculated the PLAM for the axial location when the axial location of the point source is estimated using one “summed-pixel” from each detector, obtained for each detector by adding all the pixels comprising the given sub-ROI. This Summed-Pixel Based PLAM (SPB-PLAM) represents the best axial localization accuracy that can potentially be attained by a method that utilizes only the single intensity value from each detector. Figure 1 shows plots of such SPB-PLAM values as a function of the axial position of a simulated point source when it is imaged by two-plane MUM configurations with increasing focal plane separations. For reference, the standard axial localization PLAM value for the same point source, when it is simulated as being captured in a pair of 13 × 13-pixel ROIs, is also plotted. This standard PLAM value represents the best accuracy that can potentially be achieved with an estimation algorithm like MUMLA that uses all pixel intensities as they are in both ROIs given the simulation parameters used [9]. We observe that for the 0.4 *μ*m and 0.6 *μ*m plane separations, the SPB-PLAM curves for summed-pixels of sizes 16-*μ*m and 48-*μ*m stay close to the standard PLAM curves over a large axial range of 1.5 *μ*m, indicating that for small plane separations a single-intensity-based axial localization method may suffice for some applications. However, the same is not observed for a focal plane separation of 1 *μ*m or higher, in which case no single SPB-PLAM curve stays consistently close to the standard PLAM over the same large axial range.

**Fig. 1.**
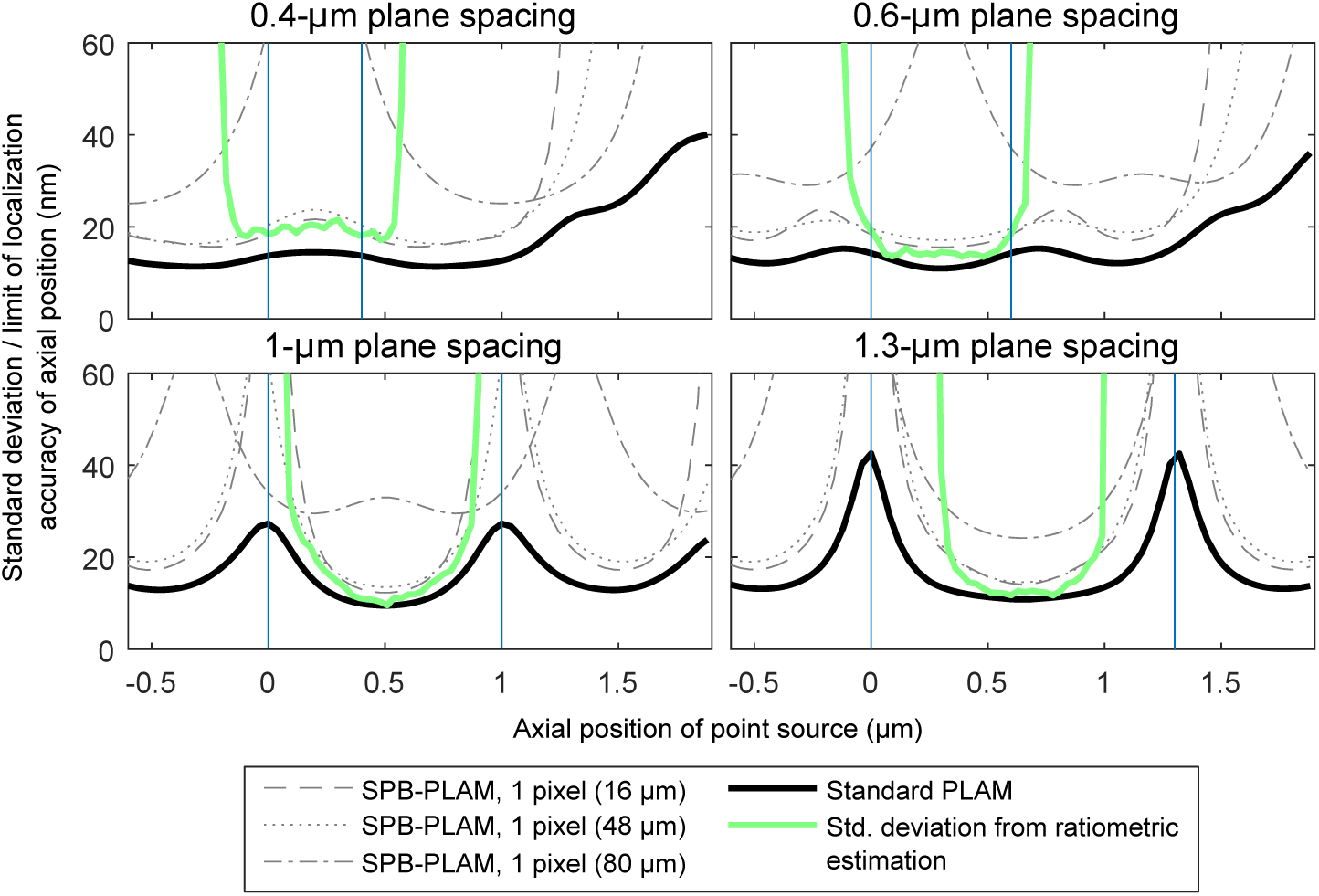
Plots of the standard PLAM and SPB-PLAMs for the axial location of a simulated point source as a function of its axial location. The standard deviation of 1000 axial location estimates obtained using the ratiometric method is also shown. Sub-panels show plots of these values in simulations with different focal plane spacings in a two-plane MUM configuration. PLAM calculations were performed with the following numerical values: magnification *M* = 100, numerical aperture *n_a_* = 1.45, pixel size: 16 *μ*m × 16 *μ*m, ROI size: 13 × 13 pixels. Here, the 16-*μ*m, 48-*μ*m and 80-*μ*m sizes of the summed-pixels correspond to areas that cover sub-ROIs of 1 × 1, 3 × 3 and 5 × 5 pixels, respectively, in the regular pixelated detectors. Calibration data for the ratiometric method was generated using data from one simulated point source with the same parameters. Standard deviation values for the ratiometric method are only plotted for axial positions where the mean of the estimates does not deviate more than 50 nm from the simulated axial position. All point sources were positioned in the center of the pixel array. Vertical lines mark the positions of the focal planes in the object space.

**Fig. 2.**
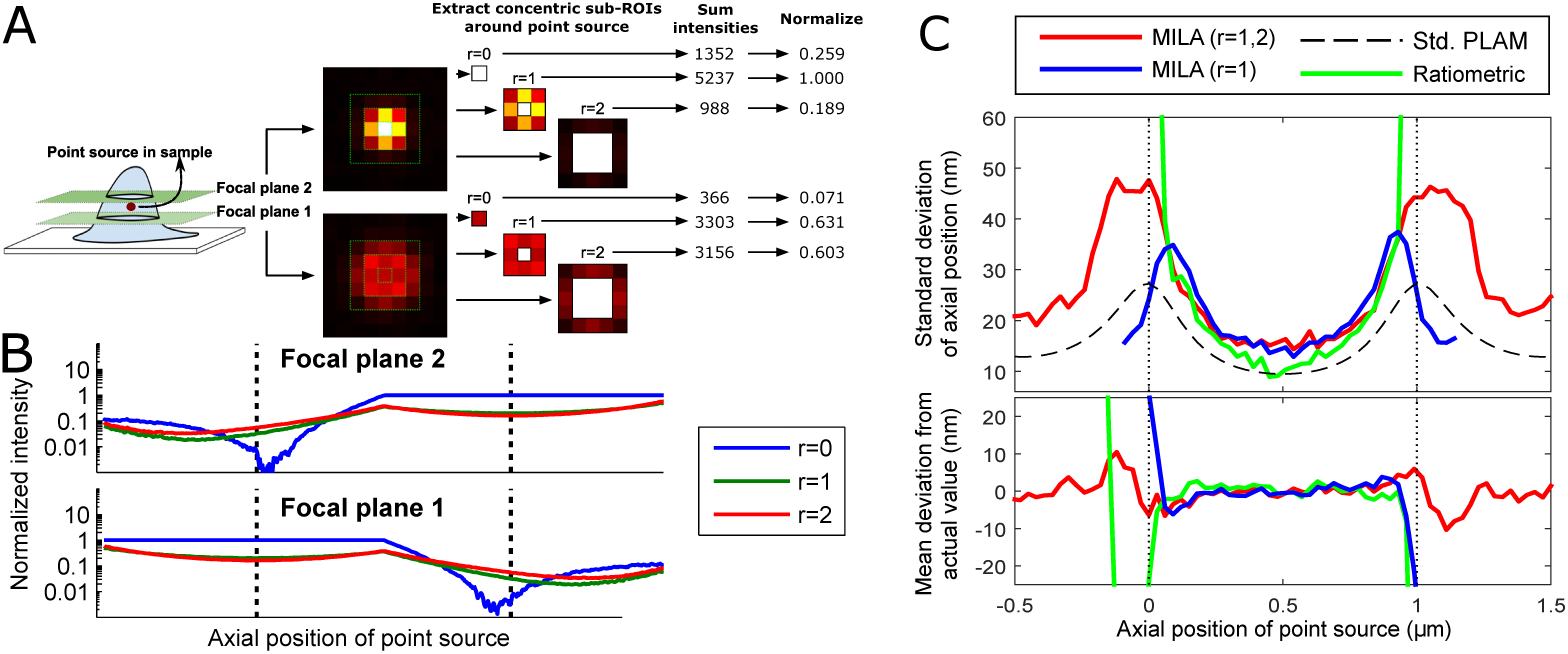
A: Schematic representation of the method of extracting multiple intensity values from images of a point source. B: Plot of normalized intensities for a typical point source as a function of its axial position. C: The top panel shows the standard deviation of axial location estimates obtained from 3 non-parametric algorithms when applied to 1000 pairs of simulated two-plane MUM images of a point source from which a mean of 4000 photons were detected per image as a function of the axial position of the point source. The bottom panel shows the deviation of the mean of these estimates from the actual axial position of the simulated point source. The red line shows the values obtained using MILA (*r* = 1, 2) and the green line represents values obtained using the ratiometric method. The blue line also shows values obtained using MILA, but using only one sub-ROI (*r* = 1). The dashed black line indicates the standard PLAM for the axial localization of the point source. The simulations for B and C were performed with the following numerical values: magnification *M* = 100, numerical aperture *n_a_* = 1.45, pixel size: 16 *μ*m × 16 *μ*m, ROI size: 13 × 13 pixels. The point source was positioned in the center of the pixel array. The calibration data was generated using data from one simulated point source with the same parameters. Vertical dotted lines mark the positions of the focal planes in the object space.

Watanabe and colleagues [10] described the ratiometric method that uses intensity estimates obtained by fitting 2D Gaussian profiles to the images of the point source from the two focal planes. The performance of this method was also tested for the same conditions using simulated images, and the standard deviation of the axial location estimates obtained by this method is shown in Fig. 1. The standard deviation was only plotted at axial locations where the mean estimated value was not separated from the simulated axial location by more than 50 nm. While the ratiometric method performs better than the SPB-PLAM values in some cases, the utility of this method is limited by the small axial region within which it is able to give estimates that are reasonably close to the true axial location.

Note that compared to all the SPB-PLAM curves, in all scenarios the standard PLAM always represents the best attainable accuracy. This is consistent with expectation because the standard PLAM applies to the case where the axial location estimator utilizes the most fine-grained spatial representation of the PSF by taking into account the individual intensity values of all the pixels that comprise the ROI.

### 3.2. Performance of MILA as a non-parametric axial location estimator for MUM data

Since the SPB-PLAM values corresponding to a single summed-pixel per detector do not remain consistently close to the standard PLAM when large focal plane spacings are used, we hypothesized that a combination of multiple summed pixels per detector might provide more axial localization information throughout the axial range of a wide-spaced MUM configuration. Hence, we propose a new method, the Multi-Intensity Lookup Algorithm (MILA), which calculates the intensity values of multiple summed pixels from the ROI in each detector. A schematic representation of the method is shown in Fig. 2A. If intensities are extracted from ROIs around the point source location as shown, we can generate a set of calibration curves (Fig. 2B) which can then be used as a lookup table. The *r* sequence specifies the number and sizes of the sub-ROIs that are chosen to be used in this method. Using just one parameter in the *r* sequence reflects a single-intensity method comparable to the previously described single intensity-based methods.

To test the performance of MILA, we simulated images of a point source located at various axial positions as acquired using a two-plane MUM configuration. We then estimated the axial location of the point source using this method with two different *r* sequences. For comparison, we also estimated the axial location using the ratiometric method and plotted the corresponding standard PLAM values (Fig. 2C). We observe that MILA (*r* = 1, 2, red line) is able to give us estimates with low standard deviation over a wide range of axial locations that even covers regions outside the two focal planes. While the ratiometric method (Fig. 2C, green line) also gives accurate estimation results for point sources situated between the two focal planes, the estimates from this method are not reliable when the point source is outside the focal planes due to bias in the estimated values (Fig. 2C, bottom panel). Thus MILA can produce consistently good localization accuracy with relatively little bias over a wider axial range than the ratiometric method. Note that when only one sub-ROI (e.g., *r* = 1, Fig. 2C, blue line) is used for intensity calculation, the results of MILA become comparable to the ratiometric method, giving high accuracies between the focal planes but being unusable outside that axial region due to unacceptably large bias.

In order to further understand the performance of MILA, we calculated a SPB-PLAM curve that gives the best accuracy attainable given the information supplied to this method. Figure 3 shows that the SPB-PLAM curve calculated with two sub-ROIs (*r* = 1, 2) trails the standard PLAM curve by less than 10 nm over a wide axial range. In contrast, SPB-PLAM curves calculated using just one sub-ROI do not display such consistency over the same range. Further, we also observe that the standard deviation of estimates obtained using MILA (*r* = 1, 2) trails the corresponding SPB-PLAM curve between the focal planes. However, this standard deviation diverges from the SPB-PLAM curve outside the focal planes, demonstrating that while MILA provides accurate axial location estimates over the entire axial range, the simple lookup method does not behave as a perfect estimation algorithm, whose accuracy would be expected to reach the corresponding SPB-PLAM.

**Fig. 3.**
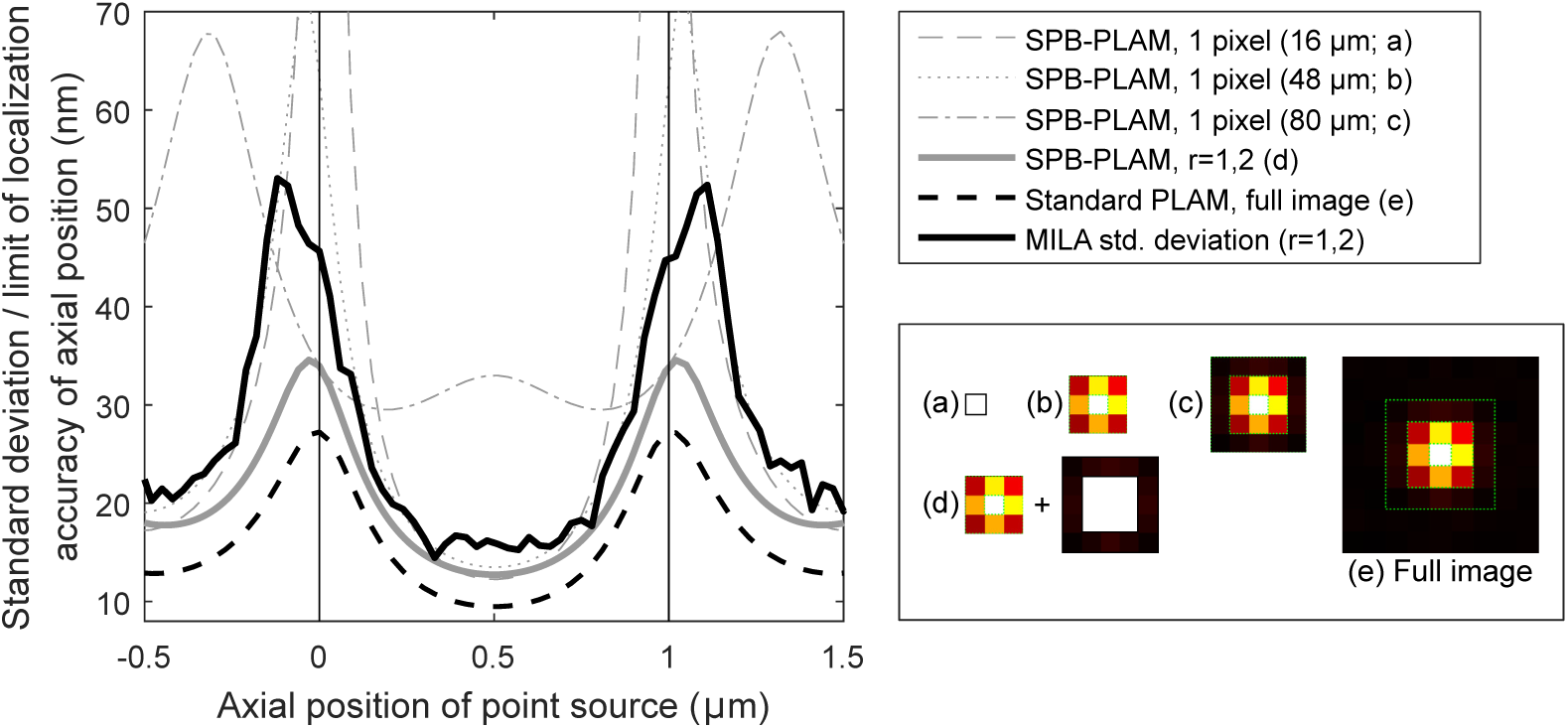
Plots of the SPB-PLAMs, the standard PLAM and the standard deviation of MILA estimates for the axial location of a simulated point source. The legend shows an example image of a point source and the pixels from that image that are summed for the different SPB-PLAM calculations. SPB-PLAM was calculated for *r* = 1, 2 by adding the Fisher information calculated for two summed-pixels whose profiles are shown in the legend. Standard deviation of axial estimates calculated using MILA (*r* = 1, 2) are also plotted. Vertical lines mark the positions of the focal planes in the object space.

### 3.3. MILA can also be used in MUM configurations with more than two focal planes

Next, we characterized the performance of MILA when applied to data from MUM configurations having more than two focal planes. One of the primary applications of MUM is to study the dynamics of single particles that traverse the entire volume of mammalian cells, which are typically 6 to 8 *μ*m thick. A four-camera MUM configuration with focal plane separations of about 1.5 to 2 *μ*m between the detectors can cover such an axial range [7]. The schematic of such a wide-spaced MUM configuration with four focal planes and the extracted ROIs and sub-ROIs are shown in Fig. 4A. We performed estimations on simulated data arising from such a configuration, where the optical parameters were kept the same as in previous cases but the number of focal planes and the inter-plane spacing alone were increased.

**Fig. 4.**
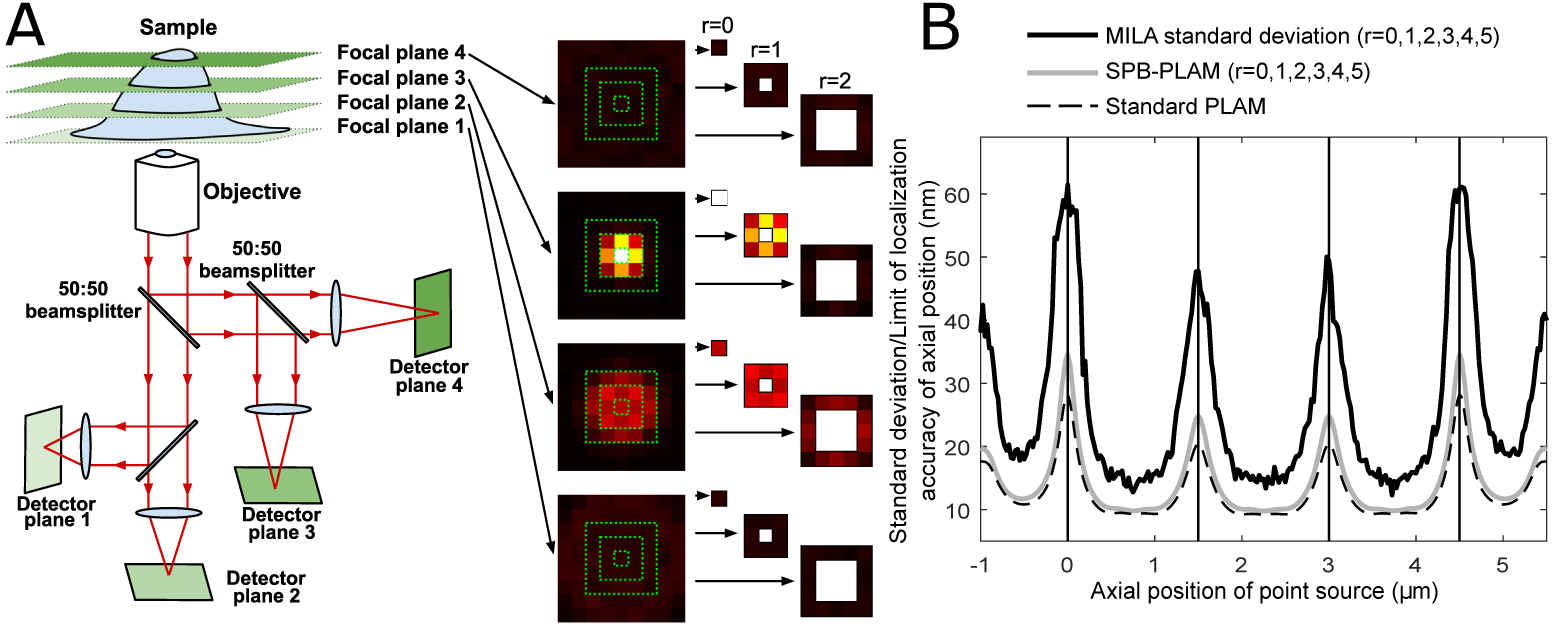
A: Schematic representation of a MUM configuration that simultaneously images four focal planes using independent detectors. Sample images of a typical point source are also shown, along with a representation of the ROIs that are used for intensity calculations. B: Plot of the accuracy reached by MILA in the estimation of the axial location of a point source using four-plane MUM data. 1000 MUM images (i.e., 1000 sets of four images corresponding to four focal planes) of a point source from which a mean of 2000 photons were detected per image were simulated and the point source's axial location was estimated using MILA. The standard deviation of these estimates is plotted as a function of the axial position of the simulated point source. The corresponding SPB-PLAM and standard PLAM for the axial location of the point source are also plotted. Simulations and estimations were performed using the following numerical values: magnification *M* = 100, numerical aperture *n_a_* = 1.45, pixel size: 16 *μ*m × 16 *μ*m, ROI size: 15 × 15 pixels, sub-ROI sequence *r* = 0, 1, 2, 3, 4, 5. The point source was positioned in the center of the pixel array. The calibration data was generated using data from one simulated point source with the same parameters. Vertical lines mark the positions of the focal planes in the object space.

As shown in Fig. 4B, even with an inter-plane spacing of 1.5 *μ*m, MILA can give axial location estimates with a standard deviation of less than 60 nm over an axial range of 6.5 *μ*m when four detectors are simulated. However, given the larger focal-plane separation, more sub-ROIs are required for MILA to perform optimally (*r* = 0, 1, 2, 3, 4, 5) and the standard deviations are consistently worse than the standard PLAM by 20 to 40 nm. Furthermore, the corresponding SPB-PLAM curve is observed to closely trail the standard PLAM, predicting that an estimator that uses the six-sub-ROI combination can produce potentially accurate axial location estimates comparable to the standard PLAM. However, the simple lookup approach used in MILA appears to be unable to perform comparably to its corresponding SPB-PLAM in such a demanding MUM configuration with wide focal plane separation. This shows that although from an information-theoretical perspective, the addition of more sub-ROIs should only increase the amount of information available to an estimator, simple lookup methods like MILA may not be able to utilize the entirety of this information in certain scenarios. Hence, with demanding wide-spaced MUM configurations, parametric estimation methods that can reach the standard PLAM may be preferred if estimating the axial location with high accuracy levels is of importance.

### 3.4. Differences in the location of the point source within the center-pixel can bias non-parametric axial location estimates

In the previous figures, the calibration data was obtained from only one point source whose position was the same as that of the test point source. However, point sources located at various positions within a single pixel will produce different images due to changes in the position of the pixel borders relative to the point source position (Fig. 5A). Hence, the intensities calculated using a non-parametric method will not be the same even for similar point sources that differ only in their sub-pixel location. When the point source generating the calibration data is not located in the same sub-pixel location as the point source for which the location is being estimated, this discrepancy can lead to bias in the estimated axial location.

**Fig. 5.**
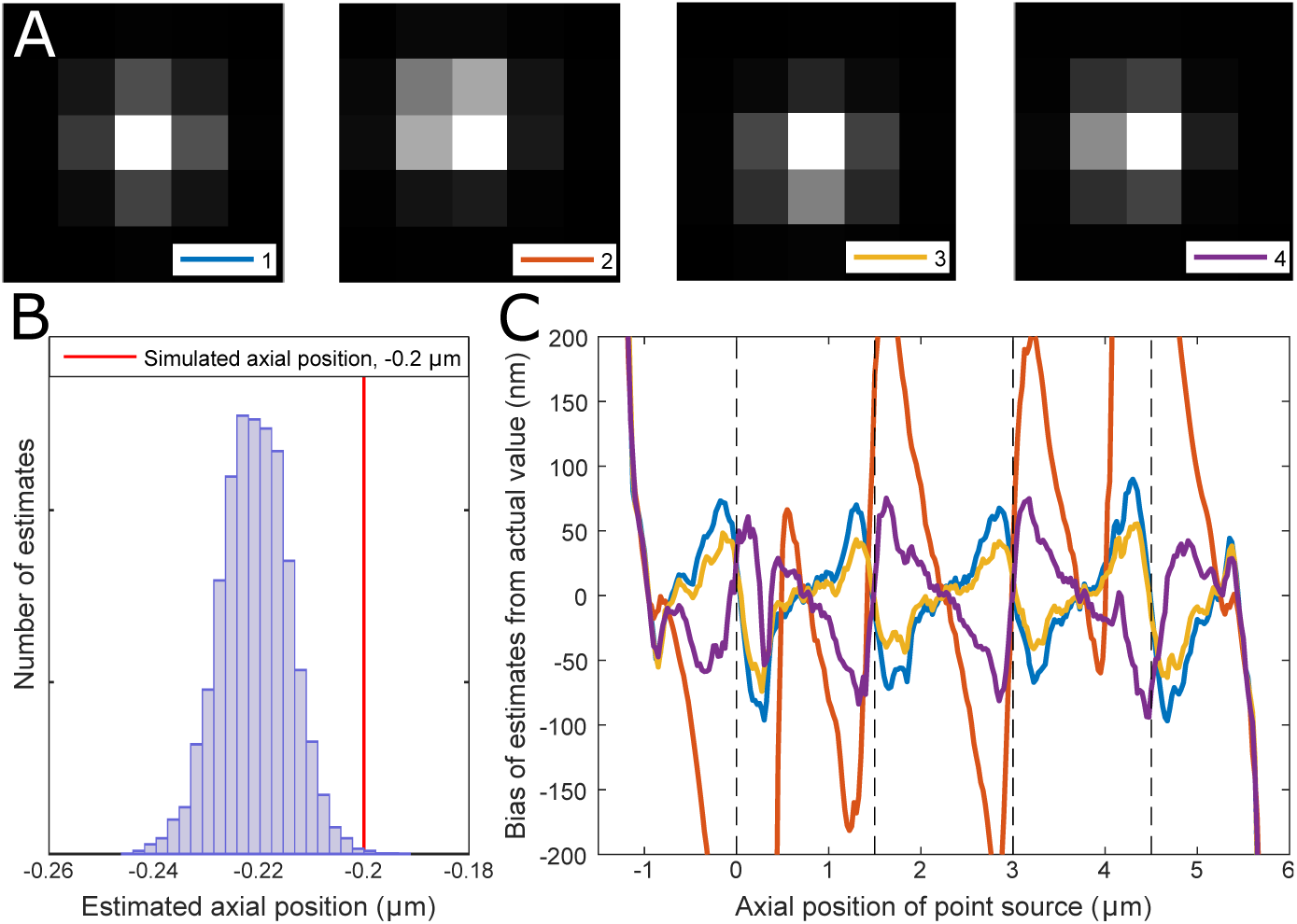
A: Example images of four simulated point sources that differ only in their locations within the center-pixel of the ROI. B: Histogram of 10000 axial location estimates of a point source obtained using MILA. Red line indicates the true axial position. C: Plot of the bias in the axial location estimates of four different point sources as a function axial location. The lateral positions of these point sources corresponds are shown in A. Each colored line represents bias in the estimates of one of the point sources. Bias is calculated as the difference between the mean of 1000 axial location estimates and the actual axial location. The axial location was estimated using MILA from simulated four-plane MUM data. Simulations and estimations were performed using the following numerical values: magnification *M* = 40, numerical aperture *n_a_* = 1.45, pixel size: 13 *μ*m × 13 *μ*m, ROI size: 11 × 11 pixels, sub-ROI sequence *r* = 0,1, 2. The calibration data was generated as a series of images of a fifth point source located at another random position within the center-pixel of the ROI and from which a mean of 10000 photons were detected per image. Vertical dotted lines mark the positions of the focal planes in the object space.

In order to demonstrate the extent of this potential bias, we simulated test images of four different point sources that only differ in their location within the center pixel of the ROI. Their axial locations were then estimated using calibration data generated using a single point source which was also located within the center pixel of the ROI. A histogram of such MILA estimates for one point source at one axial location shows a clear offset in the distribution of estimates from the simulated axial position, indicating that the deviation of the mean of these estimates might arise from systematic bias in the lookup method (Fig. 5B). When the means of such estimations are calculated at different axial positions for each of the four test point sources, we observe that there are significant deviations of these mean values from their respective simulated axial positions (Fig. 5C). We also observe that the bias curves from some point sources are more pronounced than others, indicating that the bias also appears to depend on the particular position of the point source within the pixel, with respect to the sub-pixel location of the calibration point source.

### 3.5. Including calibration data from several point sources in MILA reduces the subpixel localization bias

The bias caused in MILA by sub-pixel location differences can be reduced by using calibration data from several point sources instead of just one; this increases the probability that we have calibration data from a point source that is similar in sub-pixel position to a given point source. In order to identify the axial location of a test point source using data from several calibration point sources, we perform a global least squares minimization (Appendix 1, Eq. (3)). Figure 6A shows a plot similar to Fig. 5C, but where we utilize calibration data from 50 different point sources. The bias in the estimates has been significantly reduced at all the axial positions.

One factor that affects the quality of this bias-correction method is the number of calibration point sources to use. To study this, axial location estimations similar to those carried out for Fig. 6A were performed for two magnification-pixel size combinations using varying numbers of calibration point sources. Plotting the mean modulus bias in the axial estimates against the number of calibration point sources shows that the average bias decreases rapidly as the number of calibration point sources used increases, but does not change significantly as long as more than five calibration point sources are used (Fig. 6B).

**Fig. 6.**
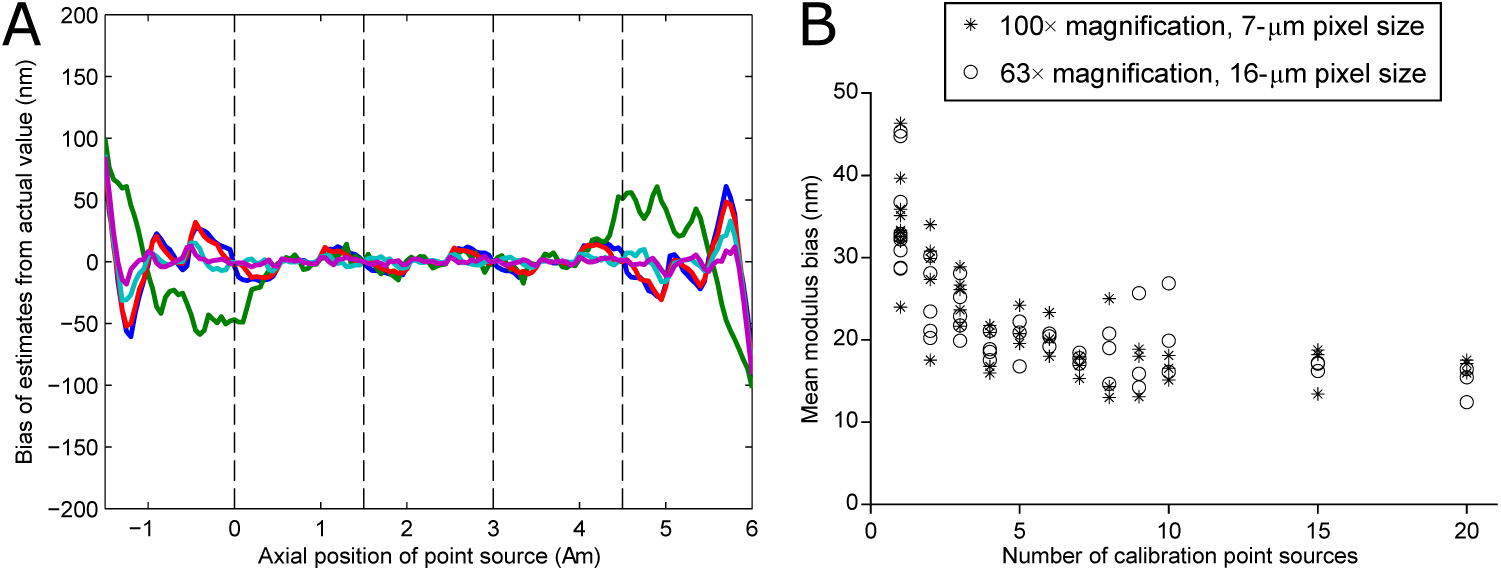
A: Plots of the difference between the mean of 1000 axial location estimates and the true axial location as a function of the axial location for five simulated point sources that are imaged using a four-plane MUM setup. Calibration data from 50 different randomly positioned point sources were used. Each colored line represents bias in the estimates of one of the point sources. Simulations and estimations were performed using the following numerical values: focal-plane separation: 1.5 *μm*, magnification *M* = 40, numerical aperture *n_a_* = 1.45, pixel size: 13 *μ*m × 13 *μ*m, ROI size: 11 × 11 pixels, sub-ROI sequence *r* = 0, 1, 2. The point sources were positioned in various locations within the center-pixel and the calibration data was generated as a series of images of 50 different point sources located in random locations within the same center-pixel and from which a mean of 10000 photons were detected per image. B: Plot of the mean modulus bias in the calculated axial location estimates as a function of the number of calibration point sources used. Test and calibration data were simulated similar to A, but calibration curves were generated with different numbers of point sources whose positions within the center-pixel were randomized every time. The modulus bias in axial estimates at axial positions across the focal plane range were then averaged and plotted against the number of calibration point sources used. All simulation parameters were maintained as above, except the magnification and pixel size were varied as indicated in the legend.

### 3.6. Evaluation of MILA using experimental data

We tested the efficacy of MILA on bead data acquired from a four-plane MUM configuration. As shown in Fig. 7A, the standard deviation for the axial location estimation of a typical bead is consistently maintained below 40 nm over a range of 4 *μ*m between the four focal planes, showing that the method can also be successfully used with experimental data. The bias in the estimated value is also kept low in these estimations (Fig. 7B).

**Fig. 7.**
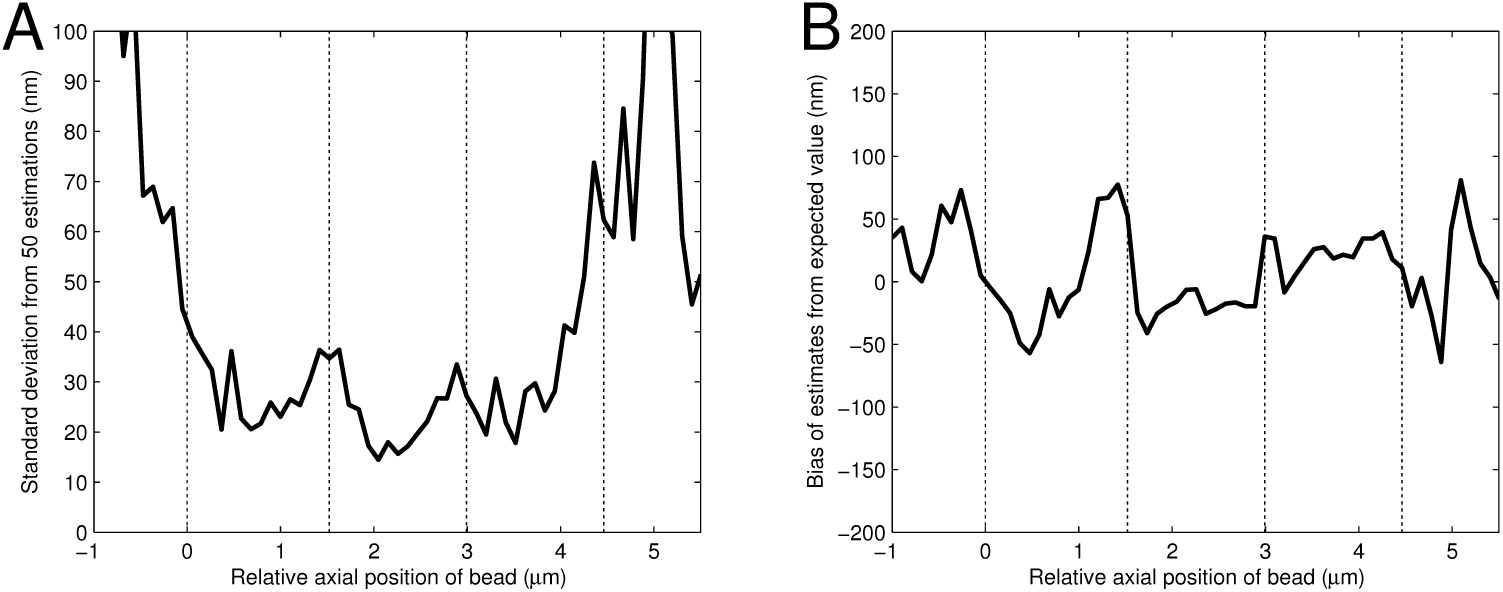
A: Standard deviation of z estimations obtained using MILA from 50 sets of four images (each corresponding to a distinct focal plane) per axial position of a typical 100nm TetraSpeck fluorescent bead as a function of the relative axial position of the bead. B: Bias of the mean estimated z value from the expected z value for the same bead as a function of its relative axial position. The following estimation parameter values were used: ROI size: 13 × 13 pixels, sub-ROI sequence *r* = 2, 3, 4. Calibration data and test data were both generated using beads from the same sample but in different fields of view. For calibration data, 10 images of the beads were acquired per detector, followed by moving the piezo nanopositioner 25 nm and repeating the process over a wide axial range. For the test data, 50 images of the beads were acquired per detector, followed by moving the piezo nanopositioner 100 nm similarly. Vertical dotted lines indicate the calculated positions of the focal planes.

## 4. Discussion

The accurate estimation of the 3D location of a point source represents a fundamental problem in single molecule microscopy with significant implications for single particle tracking applications and super-resolution microscopy. MUM has provided a solution to this 3D localization problem with the use of multiple detectors that image different focal planes and rigorous estimation algorithms like MUMLA. However, because of the relatively long processing times required by algorithms like MUMLA, a faster, easier-to-implement method may be desired. Others have implemented several related non-parametric axial localization methods which roughly abstract to comparing the relative intensities of the point source image within a small region in each detector of the MUM configuration to a lookup table.

Here we explore how much axial location information can be obtained with such intensity comparison methods by using SPB-PLAM calculations. Our observations indicate that for MUM configurations with closely-spaced focal planes (plane spacing of 0.5 *μ*m), a single intensity calculated with the correct sub-ROI size can indeed provide accurate axial location estimates. However, this phenomenon becomes less reliable with wider focal plane spacings, wherein a particular sub-ROI size may provide only accurate estimates for certain axial locations of the point source. Given the practical importance of using wide-spaced MUM configurations, an ideal non-parametric axial localization method needs to provide consistently accurate axial location estimates for point sources over a wide axial range.

Here we introduce a simple yet robust non-parametric axial location estimation method, MILA, that satisfies this important criterion. MILA also calculates the axial location from intensity estimates, but uses more than one intensity value from each focal plane image of the point source. Using simulations, we show that the accuracy attainable with this method is reasonably close to the standard PLAM over a wide axial range (Fig. 2).

One of the primary benefits of this method is its simplicity: the implementation of this method does not require any a priori information about the PSF of the microscope configuration and involves only arithmetic manipulation of the raw data that can be implemented using most software frameworks. This enables both easier implementation of the algorithm and fast computation times, which are often not possible when using parametric fitting algorithms like MUMLA.

Because we do not need to define a PSF to use MILA, it should also work with data from configurations implementing various PSF engineering methods, though highly asymmetric PSFs might attain better accuracies if we use ROI shapes that can take advantage of the intensity characteristics of those PSFs.

Further, we often observe that for the correct choice of sub-ROI combinations, the MILA estimation accuracy is reasonably close to the standard PLAM (Fig. 3), indicating that the accuracy attained by this method might be sufficient for studies where a small compromise in axial localization accuracy is permissible. However, it is also clear that results from MILA and other non-parametric methods do not fully reach the practical accuracy limits as has been shown for parametric axial location estimation methods [2].

As an example, when MILA was applied to a four-plane MUM configuration with focal planes 1.5 *μ*m apart (Fig. 4), we observed that using just a few sub-ROIs was not sufficient to obtain good estimates throughout the axial range, since the photons from the point source were spread over a much larger region in the out-of-focus detectors compared to MUM configurations with smaller focal-plane separations. Hence, intensity-based axial location estimation methods need to integrate photon counts from a larger number of pixels, increasing the added noise amounts and consequently the error in the estimation process. This phenomenon leads to the MILA standard deviation in this configuration being further separated from its corresponding SPB-PLAM curve compared to the closely spaced MUM configurations shown before (Fig. 2). It should be noted that the previously published single-intensity methods will also fail to provide reliable estimates in this configuration, showing that under such wide-spaced MUM configurations, rigorous parametric methods such as MUMLA are still the best choice for producing accurate axial location estimates. However, estimates obtained from MILA can be fed as initial conditions to such accurate parametric estimation algorithms, hence these non-parametric methods are still valuable in such cases.

One issue that affects non-parametric methods is the bias generated from the differences in the sub-pixel locations of point sources in the calibration and test data. We were able to address this problem by using lookup data from multiple calibration point sources. However, this should not cause any practical problems, since the calibration data is often obtained using fluorescent beads and utilizing data from images of many beads in a single field of view should suffice to address this sub-pixel localization bias.

Finally, we have also tested the applicability of MILA to experimental data and shown that it can accurately recapitulate the axial location of fluorescent beads. Using a four-plane MUM configuration, we were able to calculate the axial position of the beads over a range of several microns, showing the extensibility of this method to experimental data. We hope that the simplicity and versatility of this method will allow the rapid calculation of axial location estimates of point sources and make it a useful tool in applications requiring the determination of the 3D location of objects of interest from MUM data.

## APPENDIX 1: Estimating the axial location of a point source using MILA

Let us consider data collected by *D* pixelated detectors represented by *d* = 1,…, *D*, each capturing the image of a different focal plane in the object space. The images from all *D* detectors are assumed to be aligned so that the lateral positions of each pixel correspond between all the images.

The image obtained from each detector is cropped to a small ROI so that the point source of interest is situated in the center-pixel of the image. The size of the ROI is chosen such that it includes all the pixels that receive the majority of the photons from the point source. For each detector *d* = 1,…, *D*, a background estimate *B_d_* can be calculated through an appropriate method, such as taking the median of the edge pixels of the ROI or by using adjacent time-series images in super-resolution datasets.

We then calculate a series of intensity values from each ROI by summing the intensities from pixels belonging to various sub-ROIs. The shapes of the sub-ROIs chosen in this report are concentric square regions as shown in Fig. 2A. However, the shapes can be customized so that the regions of the PSF which exhibit maximal changes in intensity as a function of the point source's axial position are optimally represented in the sub-ROIs. The concentric sub-ROIs used here are represented with the sequence *r* = *r*_1_,…*r_P_*, where the sub-ROI *r_p_*, *p* = 1,…, *P* comprises all the pixels that are located less than *r_p_* pixels away from the center pixel in both the *x* and *y* dimensions but are not included as part of any smaller sub-ROI in the sequence. For example, the sequence *r* = 1, 2 represents two sub-ROIs, the 3 × 3 square region around the center pixel and the next line of pixels around this sub-ROI as depicted for scenario (d) in Fig. 3.

If *A'_d,r_p__* denotes the sum of pixel intensities for sub-ROI *r_p_* from the image of detector *d*, we normalize it as,

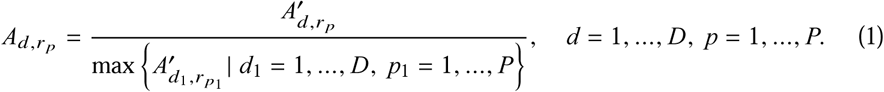

In order to calculate the axial location of the point source using the normalized intensity values, we establish a lookup table by utilizing data acquired from a calibration experiment. A calibration experiment can be defined as one where we know the axial position of the point source whose images are being acquired. In the case of simulations, we can simply simulate the images of point sources at various axial locations. In the case of practical experiments, the data can be acquired by recording images of a point source while linearly changing the focus of the microscope objective using a piezo nanopositioner. Thus we acquire *J* sets of images per detector *d*, each set corresponding to a unique axial position *z* of the piezo nanopositioner.

The images for a point source in this calibration data can then be represented by 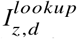, *z* = *z*_1_,…, *z_j_*, *d* = 1,…, *D*. We then calculate the sets of normalized intensity values for each lookup position to yield 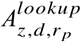, *z* = *z*_1_,…, *z_J_*, *d* = 1,…, *D*, *p* = 1,…, *P*.

Now, given a set of images *I_d_*, *d* = 1,…, *D* for an unknown point source, we can calculate an estimate *ẑ* of the point source's axial location by calculating the corresponding *A_d,r_p__* values and evaluating them against the lookup table by identifying the value of *z* for which the *A_d,r_p__* values are most similar to the 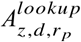 values:

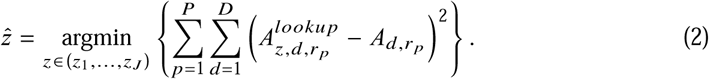

The accuracy attainable with this lookup method is, however, limited by the number of *z* values in the lookup table (for example, if we only have a lookup value every 25 nm, then the estimated axial positions can only be calculated in relatively coarse 25-nm increments). This can be addressed by calculating the 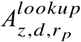 values for more *z* positions between *z_j_* and *z*_*j*+1_, *j* = 1,…, *J* − 1 by (linear) interpolation methods. Depending on the quality of the calibration data, smoothing methods may also be used to remove deviations in the calibration curve due to local variations.

In order to address sub-pixel localization bias issues, we can utilize calibration data from multiple point sources. If we have calibration data from *C* point sources, then we can represent lookup tables from all these point sources as 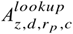, *z* = *z*_1_,…, *z_J_*, *d* = 1,…, *D*, *p* = 1,…, *P*, *c* = 1,…, *C*. Hence, when given intensities *A_d,r_p__* from an unknown point source, we globally compare them to all 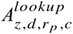 values just as above and identify the value of *z* and across lookup positions from all the calibration beads in the series represented below:

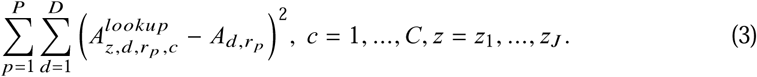

The *z* value where the above sum reaches minimum across all *c* values is then taken as the best estimate for axial location of our unknown point source, *ẑ*.

## Acknowledgments

This work was supported in part by the National Institutes of Health (R01 GM85575).

## References and links

1. P. Prabhat, S. Ram, E. S. Ward, and R. J. Ober, “Simultaneous imaging of different focal planes in fluorescence microscopy for the study of cellular dynamics in three dimensions,” IEEE Trans Nanobioscience 3, 237–242 (2004).

2. S. Ram, P. Prabhat, J. Chao, E. S. Ward, and R. J. Ober, “High accuracy 3D quantum dot tracking with multifocal plane microscopy for the study of fast intracellular dynamics in live cells,” Biophys. J. 95, 6025–6043 (2008).

3. P. A. Dalgarno, H. I. Dalgarno, A. Putoud, R. Lambert, L. Paterson, D. C. Logan, D. P. Towers, R. J. Warburton, and A. H. Greenaway, “Multiplane imaging and three dimensional nanoscale particle tracking in biological microscopy,” Opt Express 18, 877–884 (2010).

4. H. P. Kao and A. S. Verkman, “Tracking of single fluorescent particles in three dimensions: use of cylindrical optics to encode particle position,” Biophys. J. 67, 1291–1300 (1994).

5. S. R. Pavani, M. A. Thompson, J. S. Biteen, S. J. Lord, N. Liu, R. J. Twieg, R. Piestun, and W. E. Moerner, “Threedimensional, single-molecule fluorescence imaging beyond the diffraction limit by using a double-helix point spread function,” Proc. Natl. Acad. Sci. U.S.A. 106, 2995–2999 (2009).

6. M. D. Lew, S. F. Lee, M. Badieirostami, and W. E. Moerner, “Corkscrew point spread function for far-field three-dimensional nanoscale localization of pointlike objects,” Opt Lett 36, 202–204 (2011).

7. S. Ram, D. Kim, R. J. Ober, and E. S. Ward, “3D single molecule tracking with multifocal plane microscopy reveals rapid intercellular transferrin transport at epithelial cell barriers,” Biophys. J. 103, 1594–1603 (2012).

8. S. Ram, J. Chao, P. Prabhat, E. Ward, and R. Ober, “A novel approach to determining the three-dimensional location of microscopic objects with applications to 3D particle tracking,” Proc. SPIE 6443, 64430D–1 (2007).

9. A. V. Abraham, S. Ram, J. Chao, E. S. Ward, and R. J. Ober, “Quantitative study of single molecule location estimation techniques,” Opt Express 17, 23352–23373 (2009).

10. T. M. Watanabe, T. Sato, K. Gonda, and H. Higuchi, “Three-dimensional nanometry of vesicle transport in living cells using dual-focus imaging optics,” Biochem. Biophys. Res. Commun. 359, 1–7 (2007).

11. H. I. Dalgarno, P. A. Dalgarno, A. C. Dada, C. E. Towers, G. J. Gibson, R. M. Parton, I. Davis, R. J. Warburton, and A. H. Greenaway, “Nanometric depth resolution from multi-focal images in microscopy,” J R Soc Interface 8, 942–951 (2011).

12. M. Born and E. Wolf, Principles of Optics: Electromagnetic Theory of Propagation, Interference and Diffraction of Light (Cambridge University Press, 1999).

13. “EstimationTool,” http://wardoberlab.com/software/estimationtool.

14. “FandPLimitTool,” http://wardoberlab.com/software/fandplimittool.

